# A multi-modal image fusion workflow incorporating MALDI imaging mass spectrometry and microscopy for the study of small pharmaceutical compounds

**DOI:** 10.1101/2024.03.12.584673

**Authors:** Zhongling Liang, Yingchan Guo, Abhisheak Sharma, Christopher R. McCurdy, Boone M. Prentice

**Author notes:** Address correspondence to: Dr. Boone M. Prentice 214 Leigh Hall, PO Box 117200, Department of Chemistry University of Florida Gainesville, FL 32611, USA.

## Abstract

Multi-modal imaging analyses of dosed tissue samples can provide more comprehensive insight into the effects of a therapeutically active compound on a target tissue compared to single-modal imaging. For example, simultaneous spatial mapping of pharmaceutical compounds and endogenous macromolecule receptors is difficult to achieve in a single imaging experiment. Herein, we present a multi-modal workflow combining imaging mass spectrometry with immunohistochemistry (IHC) fluorescence imaging and brightfield microscopy imaging. Imaging mass spectrometry enables direct mapping of pharmaceutical compounds and metabolites, IHC fluorescence imaging can visualize large proteins, and brightfield microscopy imaging provides tissue morphology information. Single-cell resolution images are generally difficult to acquire using imaging mass spectrometry, but are readily acquired with IHC fluorescence and brightfield microscopy imaging. Spatial sharpening of mass spectrometry images would thus allow for higher fidelity co-registration with higher resolution microscopy images. Imaging mass spectrometry spatial resolution can be predicted to a finer value via a computational image fusion workflow, which models the relationship between the intensity values in the mass spectrometry image and the features of a high spatial resolution microscopy image. As a proof of concept, our multi-modal workflow was applied to brain tissue extracted from a Sprague Dawley rat dosed with a kratom alkaloid, corynantheidine. Four candidate mathematical models including linear regression, partial least squares regression (PLS), random forest regression, and two-dimensional convolutional neural network (2-D CNN), were tested. The random forest and 2-D CNN models most accurately predicted the intensity values at each pixel as well as the overall patterns of the mass spectrometry images, while also providing the best spatial resolution enhancements. Herein, image fusion enabled predicted mass spectrometry images of corynantheidine, GABA, and glutamine to approximately 2.5 μm spatial resolutions, a significant improvement compared to the original images acquired at 25 μm spatial resolution. The predicted mass spectrometry images were then co-registered with an H&E image and IHC fluorescence image of the μ- opioid receptor to assess co-localization of corynantheidine with brain cells. Our study also provides insight into the different evaluation parameters to consider when utilizing image fusion for biological applications.

## INTRODUCTION

The study of pharmacology, toxicity, and localization of a drug in target tissues is an important area of research in pharmaceutical sciences.^1–3^ Imaging analyses of dosed tissue samples can provide insight into the spatial domain, where both morphological and molecular information can be acquired depending on the imaging technique. Morphological imaging techniques allow for the visualization of different structures in the tissue and molecular imaging techniques provide biochemical information such as metabolite and protein expression.^4^ A wide variety of imaging techniques have been applied to areas of research such as studying pharmacokinetics, assessing drug efficacy, and mapping biomarkers.^1,5^ For example, imaging mass spectrometry enables the label-free mapping of endogenous and exogenous compounds and has emerged as a useful tool for studying drug pharmacology.^6,7^ Unlike techniques such as autoradiography and positron emission tomography (PET), imaging mass spectrometry does not require radiolabeling and can readily distinguish between a parent compound and its metabolites.^8,9^ We have recently used imaging mass spectrometry to map the brain spatial distributions of alkaloids derived from *Mitragyna speciosa*, a species of plant native to Southeast Asia commonly known as kratom and used to reduce fatigue, treat pain, and treat opioid withdrawal symptoms.^10–13^ Although it is known that the alkaloid components of kratom are the major contributors to its therapeutic effects, the specific alkaloids and mechanisms of action remain unclear. Kratom is currently not approved by the U.S. Food and Drug Administration (FDA) because of these potential health risks and the unknown effects on brain chemistry, highlighting the need for increased study by imaging technologies.^14–17^

A single imaging modality is often unable to provide a comprehensive view of the total morphological and molecular information desired. For example, morphological imaging techniques do not provide the localization of specific biological molecules, while molecular imaging techniques may not provide sufficient spatial or morphological information. Thus multi-modal approaches, combining two or more different imaging modalities, can offer a more holistic view of the tissue.^18^ In this study, a multi-modal imaging workflow incorporating brightfield microscopy, immunohistochemistry (IHC), and matrix-assisted laser desorption/ionization (MALDI) imaging mass spectrometry was developed. Brightfield microscopy of hematoxylin and eosin (H&E)-stained tissue sections is routinely used to assess tissue morphology.^19^ IHC is commonly used to map proteins and receptors of interest to assess potential pharmaceutical targets and metabolism.^20^ MALDI imaging mass spectrometry is a label-free imaging technique used to map spatial distributions of metabolites, pharmaceutical compounds, lipids, and proteins of interest.^21–28^ Imaging mass spectrometry has been previously combined with other imaging modalities such as IHC, autofluorescence imaging, and IR spectroscopy in multi-modal imaging workflows.^29–33^. While imaging modalities such as brightfield microscopy and fluorescence imaging routinely achieve single cell spatial resolutions, the spatial resolution of MALDI imaging mass spectrometry is generally the most limited of these multi-modal techniques (*i.e.*, 10-200 μm).^34,35^ Enhancing the spatial resolution of the mass spectrometry image to a value closer to that of the other imaging modalities promises better data and image integration. Experimental efforts to obtain high spatial resolution imaging mass spectrometry data (*i.e.*, 1-10 μm) for improved integration with higher resolution optical and fluorescence imaging modalities come at the cost of lower sensitivity and often require specialized instrumentation.^36–40^

One alternative approach to achieving high spatial resolution imaging mass spectrometry is to computationally predict these images to finer spatial detail. This can be performed by mathematically combining the mass spectrometry images with an image obtained from a higher spatial resolution modality, a process that has been termed “image fusion.”^41^ A cross-modality mathematical regression model is constructed that allows for variables in one technology (*e.g.*, imaging mass spectrometry) to be predicted using variables from another technology (*e.g.*, microscopy). Image fusion with mass spectrometry data has been demonstrated using techniques such as brightfield microscopy and IR spectroscopy.^31,41,42^ For example, image fusion performed using lipid imaging mass spectrometry data and brightfield microscopy images enabled 10-fold spatial sharpening of the mass spectrometry images.^41^ This workflow involves two important steps: (1) alignment of the two images in the same coordinate plane and (2) modeling the cross-modality relationship between the intensity variables in the mass spectrometry images and the H&E image features. The H&E image features then serve as input data for the model to generate the high spatial resolution mass spectrometry image. Linear regression and partial least squares regression (PLS) are two commonly used mathematical models for image fusion.^41^ Deep learning models such as convolutional neural networks (CNN) have also recently been employed in the image fusion field, though much remains to be explored.^43,44^ Additionally, the use of random forest regression, a non-linear regression model, has not been explored in this type of image fusion workflow. Herein, linear regression, PLS, CNN, and random forest regression were used to test a wide variety of image fusion models.

A proof-of-concept multi-modal imaging workflow was applied to brain tissues extracted from rats dosed with corynantheidine, which is a minor alkaloid component of kratom, to provide a more holistic view of this alkaloid’s therapeutic activity in the brain. MALDI imaging mass spectrometry was used to visualize the intrabrain spatial localization of corynantheidine as well as neurotransmitter metabolites. IHC fluorescence imaging was also used to visualize μ-opioid receptors. Previous studies have demonstrated the binding affinities of kratom alkaloids toward opioid receptors, such as the partial agonist action of corynantheidine towards human μ-opioid receptors in cell-based assays.^45–47^ Multimodal image fusion models built using H&E brightfield microscopy are first used to spatially sharpen the mass spectrometry images acquired at 25 μm spatial resolution to a predicted spatial resolution of 2.5 μm. The upsampled imaging mass spectrometry data is then co-registered with IHC fluorescence imaging for identification of potential receptor targets. This workflow allows for improved alignment and fidelity of the mass spectrometry and IHC images, enabling improved insight into kratom alkaloid localization.

## RESULTS

### Imaging Mass Spectrometry

Imaging mass spectrometry analysis was performed on brain tissue extracted from Sprague Dawley rats intravenously injected with the kratom alkaloid, corynantheidine. DCTB was used as a MALDI matrix in this study to improve the alkaloid limit of detection and enable spatial resolutions of 75 μm and 25 μm (**Supplemental Figure 1 & 2**), which is much higher than the 200 μm spatial resolution employed in our previous imaging mass spectrometry analyses of kratom alkaloids.^12^ Corynantheidine is localized to the white matter, which is evident in the 25 μm spatial resolution image (**Supplemental Figure 2a**). We have previously noted that dehydrogenated [M - H]^+^ and [M - 3H]^+^ alkaloid ion types can be formed under certain MALDI analysis conditions (*e.g.*, employing DCTB as a MALDI matrix). While this can complicate spectral and imaging analyses, the improved alkaloid limit of detection enabled by DCTB here is necessary for high spatial resolution imaging and enables this proof-of-concept image fusion demonstration. In negative ion mode, a DAN MALDI matrix enables detection of GABA and glutamine on a serial tissue section, which are localized to the molecular layer (**Supplemental Figure 2b and 2c**). Kratom is known to treat pain and opioid withdrawal symptoms,^10,11^ and the alkaloidal components of kratom are the main contributors to the analgesic effects of kratom. GABA is a neurotransmitter involved in the chronic pain pathway, and glutamine is a precursor of GABA and other neurotransmitters.^48,49^ Investigating the spatial distributions of GABA and glutamine, as well as other metabolites, in rat brain tissue dosed with corynantheidine may provide insight into the pharmacological properties of corynantheidine.

### Image Fusion

Image fusion was performed to enhance the imaging mass spectrometry spatial resolution by approximately 10-fold in order to more accurately integrate the mass spectrometry images of corynantheidine, GABA, and glutamine with other imaging modalities. Briefly, mass spectrometry images, an autofluorescence image following MALDI analysis, and a subsequent H&E image were acquired from the target tissue section (**Figure 1**). The mass spectrometry image was first aligned with the autofluorescence image by matching the laser ablation marks in the autofluorescence image to the corresponding pixels in the mass spectrometry image. The autofluorescence image was then aligned to the H&E image based on the corresponding tissue morphology.^50^ Following co-registration, feature extraction to 120 features was then performed using the H&E image. Finally, mathematical modeling using one of the candidate models was performed to describe the spatial relationships between the H&E features and the intensity values of the target compound in the mass spectrometry image. The model enables predicted high spatial resolution mass spectrometry images by using the H&E image features as the input. Model optimization was individually performed for both corynantheidine and GABA mass spectrometry images. Since GABA and glutamine showed similar spatial distributions, the optimized model for GABA is also expected to perform well for glutamine. Candidate models of linear regression, partial least squares regression (PLS), random forest regression, and two-dimensional convolutional neural networks (2-D CNN) were evaluated. Linear regression and PLS regression model were the original linear models used in the first report of mass spectrometry image fusion.^41^ Random forest regression is a nonlinear regression model yet to be explored in mass spectrometry image fusion and CNN models have recently been explored as potential image fusion engines.^43,44,51^ The CNN models are attractive options for image fusion because these models only require the original RGB features of the H&E image. The CNN models are able to extract features using convolutional neural network layers, eliminating the need for manual feature extraction (**Supplemental Table 2**).^52,53^ Multiple permutations of each model were evaluated to determine the model that best captured the relationship between the intensity values in the mass spectrometry image and the H&E image features. For example, all 120 H&E image features were used for both the linear and PLS models. The component number (ranging from 1 to 120) was optimized for the PLS model. The number of decision trees (ranging from 1 to 100) and the number of H&E image features (3, 15, 30, 60, or 120) used as data input were optimized for the random forest model.

**Figure 1.**
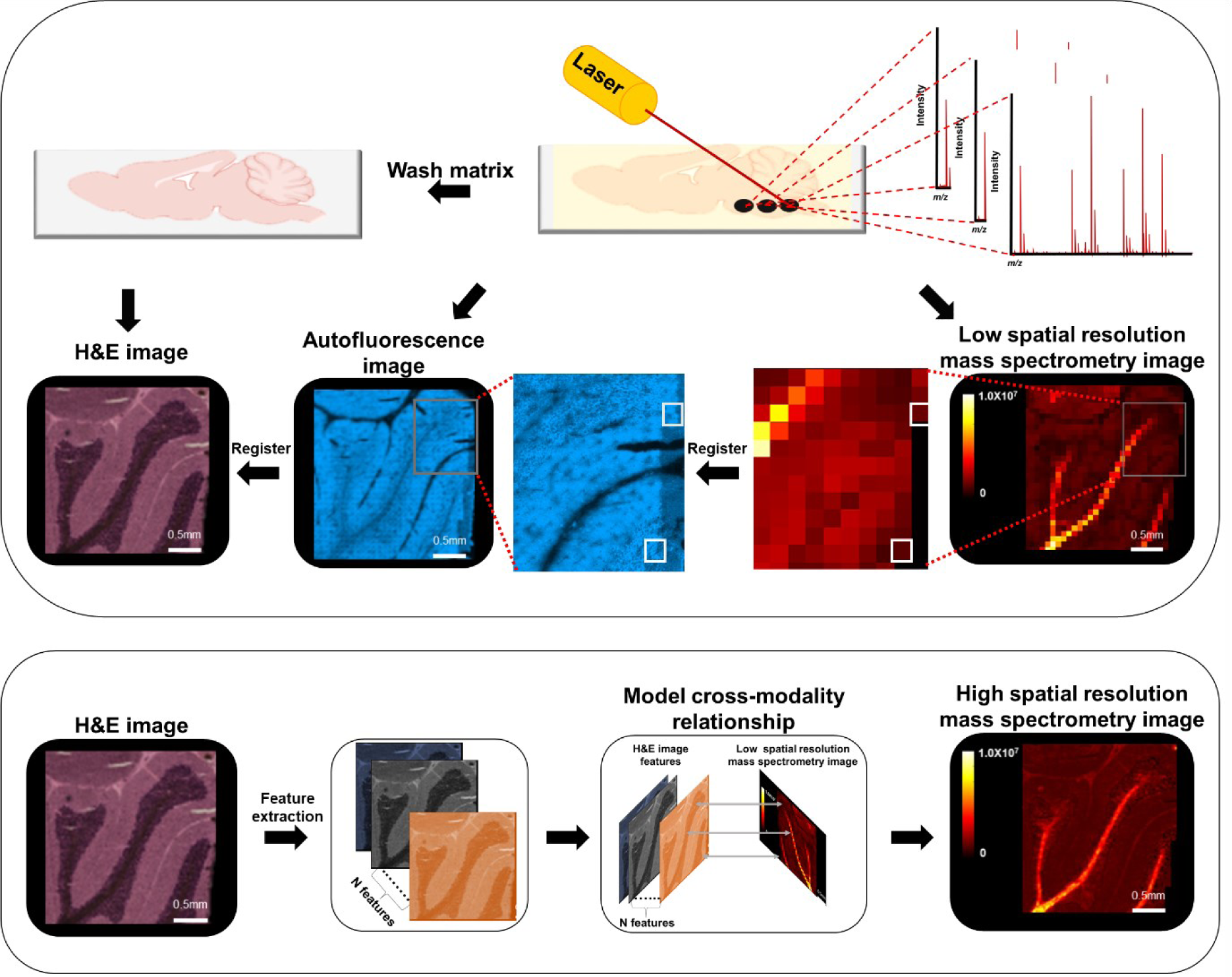
The image fusion workflow overview. The workflow consists of two main steps: (1) registration of the mass spectrometry image to the H&E image and (2) modeling the cross-modality relationship between the H&E image and the mass spectrometry image. The top portion of the figure depicts step 1, where imaging mass spectrometry is first performed on the target tissue. An autofluorescence image is acquired to visualize the laser ablation marks produced during imaging mass spectrometry. H&E staining is then performed and the slide and is imaged using brightfield microscopy. The mass spectrometry image is registered to the autofluorescence image using the corresponding laser ablation marks in the autofluorescence image and the pixels in the mass spectrometry image. The autofluorescence image is then registered to the H&E image using the tissue morphology. Magnified regions of mass spectrometry and autofluorescence images are highlighted using the gray boxes. The corresponding pixels in the autofluorescence and mass spectrometry image are highlighted using the white boxes. The bottom portion of the figure depicts step 2, where feature extraction is performed on the H&E image to acquire 120 features. The cross-modality relationship between the H&E image features and the intensity values in the mass spectrometry image is then modeled to predict the high spatial resolution mass spectrometry image.

The accuracies of the predicted mass spectrometry images were evaluated based on the relative average residual value, the correlation scores (R^2^), and the spatial resolutions of the images. The relative average residual values are calculated using the average of the absolute values of the intensity differences between the original and predicted images, divided by the average intensity of the original mass spectrometry image (see Equation 1 in Methods). A smaller relative average residual value indicates that the model is more accurately predicting the intensity values. The correlation score is the R^2^ value acquired by comparing the predicted intensity value for each pixel in the predicted mass spectrometry image to the corresponding intensity value in the original mass spectrometry image. A correlation score close to 1 implies that the model is accurately predicting the spatial pattern of the image. Different degrees of spatial sharpening can be achieved with different models, even though the output pixel number of the predicted mass spectrometry images will always be the same as the input H&E image (*i.e.*, the spatial resolution reported here of the predicted mass spectrometry image is solely based on the output pixel number). It should be noted that we have used the term “spatial resolution” herein to be consistent with prior literature, though “pixel size” is likely more accurate terminology. “Perfect” relative average residual values and correlation scores indicate that the model performs best when image fusion generates the exact same predicted image as the low spatial resolution input image. Consequently, the predicted mass spectrometry image was also magnified to evaluate the degree of spatial sharpening.

Mass spectrometry images acquired at a spatial resolution of 75 μm and H&E images acquired at a spatial resolution of roughly 7.5 μm were initially used for mathematical model optimization (**Figure 2 & Supplemental Figure 3**). Linear regression did not require any optimization, the component number was optimized for the PLS model, the decision tree number was optimized for the random forest model, and 10,000 epochs were used for the 2-D CNN model. The PLS regression model using 40 components predicted the corynantheidine mass spectrometry image with the lowest relative average residual (0.380) and highest correlation score (R^2^ = 0.791) compared to other PLS regression models that predicted images with similar levels of spatial sharpening compared to the original image (**Figure 2a** & **Supplemental Table 3**). The spatial sharpening of the predicted image compared to the original image should be visually inspected in addition to considering the relative average residual and correlation score. The random forest model using 100 decision trees and 30 H&E image input features predicted the corynantheidine mass spectrometry image with the lowest relative average residual (0.318) and highest correlation score (R^2^ = 0.834) when compared to other random forest models that predicted corynantheidine images with similar levels of spatial sharpening compared to the original image (**Supplemental Table 3**). Of the four optimized models, this random forest regression model performed best in spatial sharpening (0.318 relative average residual value, R^2^ = 0.834) (**Supplemental Figure 4a**). The 2-D CNN model (0.299 relative average residual value, R^2^ = 0.849) performed best in minimizing the relative average residual value and maximizing the correlation score (**Figure 2a**).

**Figure 2.**
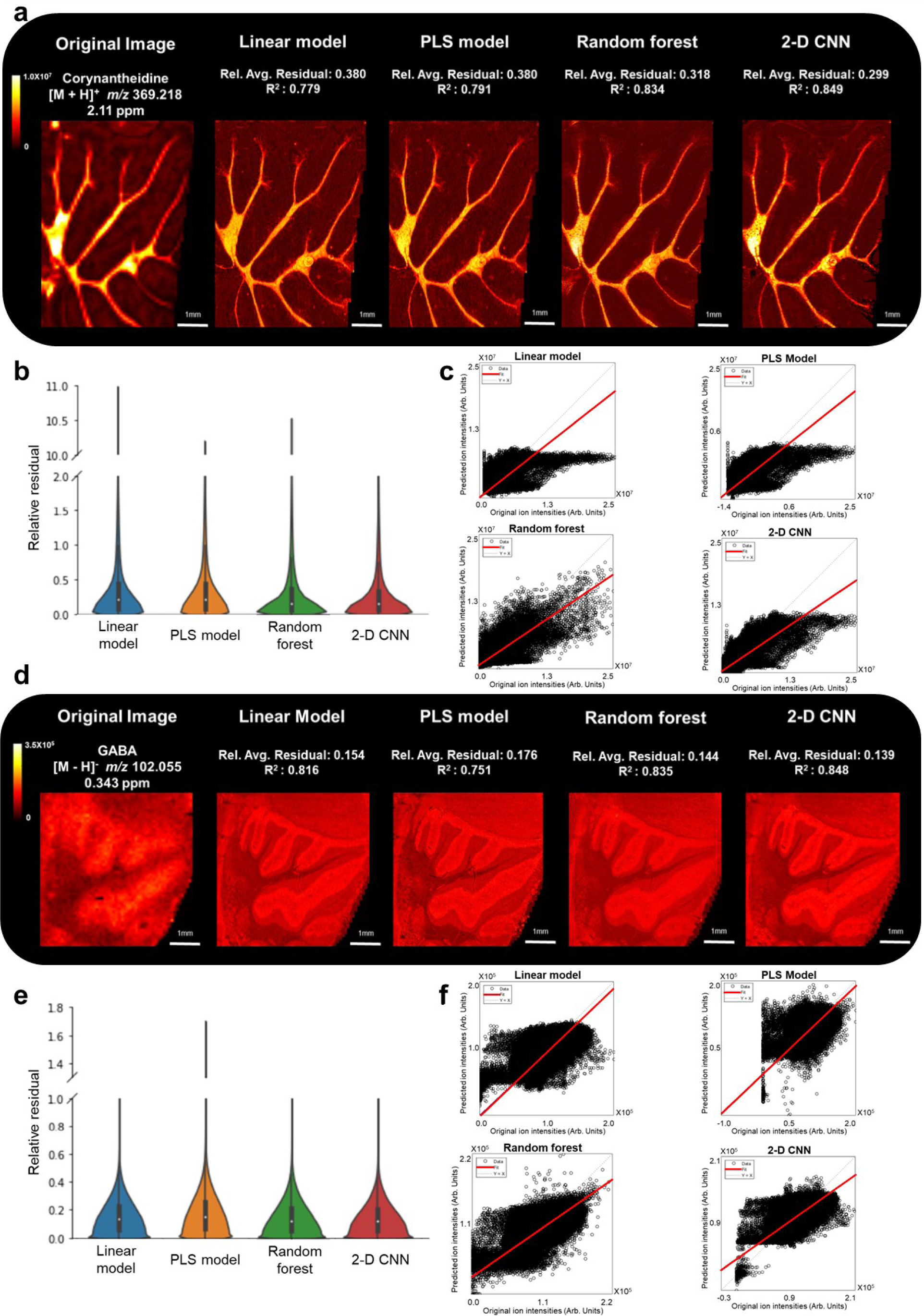
Image fusion results from 75 μm spatial resolution mass spectrometry images. Image fusion was performed on mass spectrometry images of (**a-c**) corynantheidine and (**d-f**) GABA using linear regression, PLS regression, random forest regression, and 2-D CNN. The relative average residual (Rel. Avg. Residual) and correlation score (R^2^) are calculated for each predicted mass spectrometry image. The relative residual values for each pixel, which are utilized in calculating the relative average residual, are depicted using a violin plot. The correlation score is obtained by plotting the original mass spectrometry image intensity values against the predicted mass spectrometry image intensity values.

Model optimization was also performed using the mass spectrometry image of GABA acquired at 75 μm spatial resolution (**Figure 2d**). The PLS regression model using 10 components predicted the GABA mass spectrometry image with the lowest relative average residual value (0.176) and highest correlation score (R^2^ = 0.751) compared to other PLS regression models that predicted images with similar levels of spatial sharpening compared to the original image (**Supplemental Table 4**). Similar to the random forest regression model optimization for corynantheidine, the random forest model using 100 decision trees and 30 H&E image input features was most accurate for GABA prediction. This random forest model (0.144 relative average residual value, R^2^ = 0.835) performed best in spatial sharpening (**Supplemental Figure 4b**) and the 2-D CNN model (0.139 relative average residual value, R^2^ = 0.848) again performed best in minimizing the relative average residual value and maximizing the correlation score. These same two models were used to spatially sharpen the glutamine mass spectrometry image acquired at 75 μm spatial resolution (**Supplemental Figure 3**). Indeed, the glutamine mass spectrometry images predicted using both the random forest model (0.144 relative average residual value, R^2^ = 0.841) and the 2-D CNN model (0.135 relative average residual value, R^2^ = 0.859) resulted in similar relative average residual values and correlation scores compared to the predicted GABA images.

### Multi-modal Imaging

Mass spectrometry images of corynantheidine, GABA, and glutamine were acquired at 25 μm spatial resolution for incorporation into the multi-modal imaging workflow (**Figure 3**). Image fusion performed using random forest regression (using 100 decision trees and 30 H&E image input features) and 2-D CNN models enabled approximately 10-fold spatial sharpening of the mass spectrometry images. All models demonstrated improved spatial sharpening compared to the original mass spectrometry image (**Supplemental Figure 5).** The 2-D CNN model (0.399 relative average residual value, R^2^ = 0.847) performed better for corynantheidine in minimizing the relative average residual value and maximizing the correlation score compared to the random forest regression model (0.446 relative average residual value, R^2^ = 0.807) (**Figure 3a**). The random forest regression model performed better in spatial sharpening (**Supplemental Figure 5a**). The accuracy of the prediction was prioritized for this application and thus guided selection of the 2-D CNN model for the final workflow, which still enabled resolution of cellular structures (**Figure 4** and **Supplemetal Figure 6**). The random forest regression model performed better overall for GABA and glutamine (GABA: 0.141 relative average residual value, R^2^ = 0.655; glutamine: 0.174 relative average residual value, R^2^ = 0.643) compared to the 2-D CNN model (GABA: 0.142 relative average residual value, R^2^ = 0.634; glutamine: 0.174 relative average residual value, R^2^ = 0.636) (**Figure 3d** and **3g** and **Supplemental Figure 5b** and **5c**). Surprisingly, the correlation scores for the 2.5 μm predicted mass spectrometry images of both GABA and glutamine were lower than those for the 7.5 μm predicted mass spectrometry images, while the relative average residual values are essentially the same. Strong correlation between mass spectrometry and H&E images requires that the relative abundance of the target analyte in the mass spectrometry image change with different biological morphologies that are detected in the H&E image. The higher spatial resolution 25 μm mass spectrometry image results in lower signal for GABA and glutamine analytes, diminishing the limit of detection and dynamic range. This more limited dynamic range may result in a poorer spatial correlation between the mass spectrometry and H&E images, making spatial pattern recognition more difficult and resulting in lower correlation scores. The relative average residual value is an average value across the entire image, meaning it is less affected by the intensity fluctuations of single pixels. This highlights the importance of mass spectrometry image quality in the image fusion process. Future work may explore on-tissue chemical derivatization of GABA and glutamine via 2,3-diphenyl-pyranylium tetrafluoroborate (DPP-TFB) to improve ionization efficiencies,^54,55^ thereby improving limits of detection and dynamic range for higher quality predicted fusion images. The robustness of the optimized image fusion models was further evaluated by calculating the standard deviation image. The standard deviation image is similar to the confidence interval image proposed by Van de plas.^41^ Briefly, 10 bootstrap models were generated for each random forest regression model (GABA and glutamine) and 5 bootstrap models were generated for the 2-D CNN model (corynantheidine) (**Supplemental Figure 7**). The standard deviations of the predictions by the set of bootstrap models were plotted across the tissue area to produce the standard deviation image. Areas in the image that have a high standard deviation indicate lower confidence in the predicted ion intensity values, while areas with a low standard deviation indicate higher confidence in the predicted ion intensity values (**Supplemental Figure 8**). The main purpose of the standard deviation image is to provide the user with an evaluation parameter to assess whether the prediction results are robust enough for their study (*i.e.*, in the absence of having a ground truth high spatial resolution mass spectrometry image, which is not always available).

**Figure 3.**
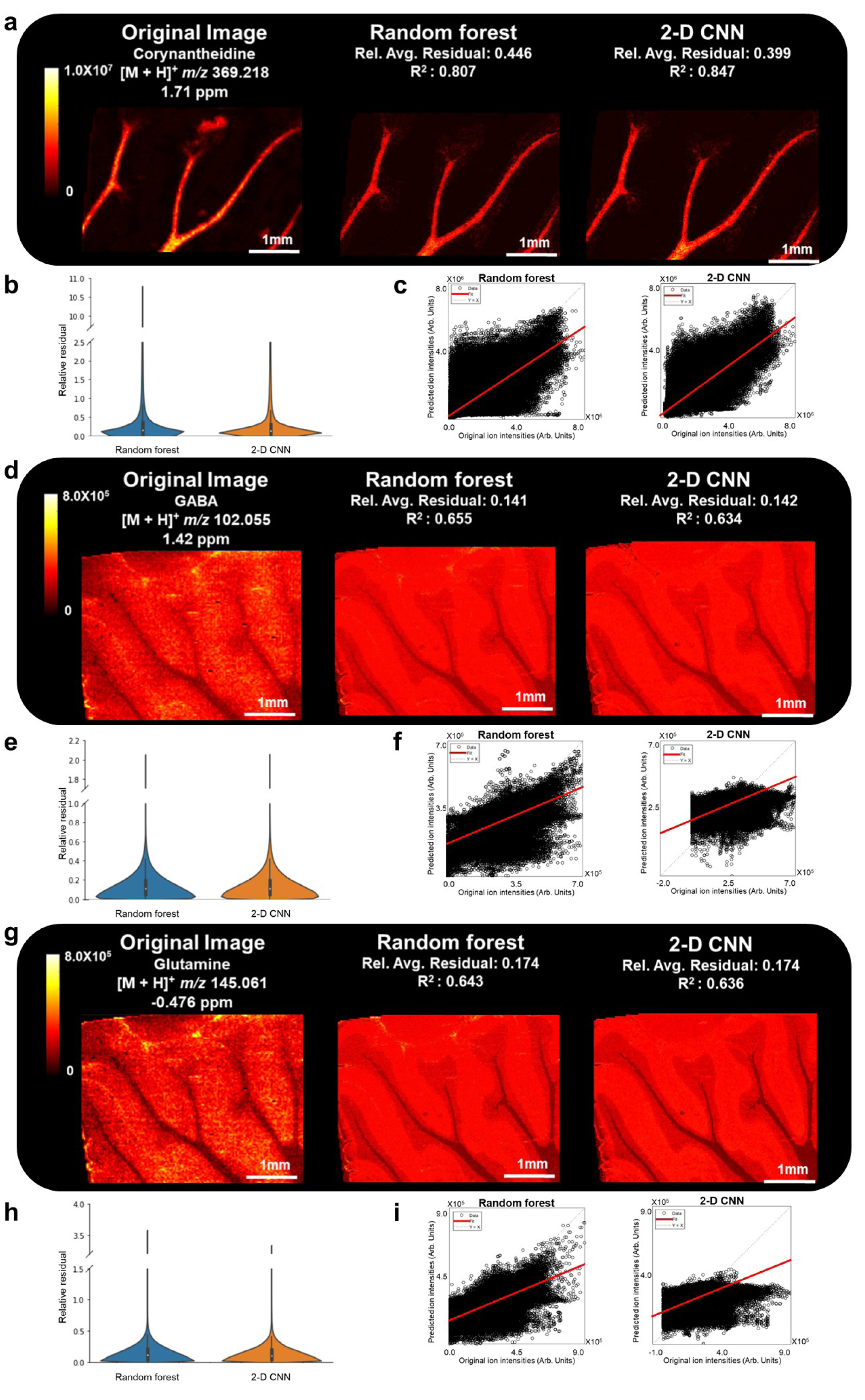
Image fusion results from 25 μm spatial resolution mass spectrometry images. Image fusion performed on mass spectrometry images of (**a-c**) corynantheidine, (**d-f**) GABA, and (**g-i**) glutamine using random forest regression and 2-D CNN models results in predicted 2.5 μm spatial resolution images. The relative average residual values (Rel. Avg. Residual) and correlation scores (R^2^) are calculated for each predicted mass spectrometry image. The relative residual values for each pixel, which are utilized in calculating the relative average residual, are depicted using a violin plot. The correlation score is obtained by plotting the original mass spectrometry image intensity values against the predicted mass spectrometry image intensity values.

**Figure 4.**
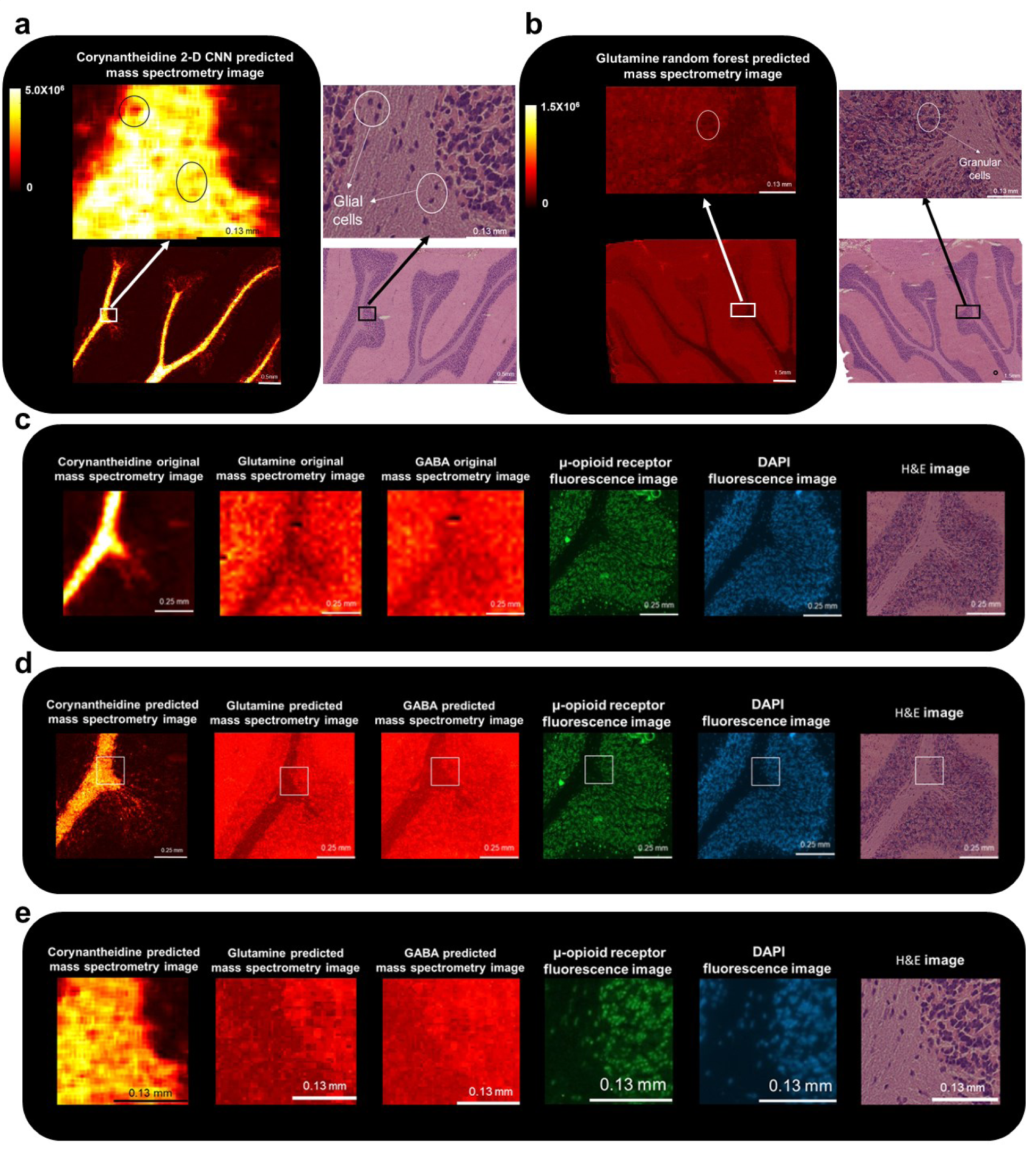
Multi-modal imaging workflow. Predicted fusion images of (**a**) corynantheidine and (**b**) glutamine are co-registered with H&E microscopy images to visualize brain morphology and cell bodies. The circles highlight corresponding glial cells (corynantheidine image) and granular cells (glutamine image) in the predicted mass spectrometry and H&E images. The final multi-modal imaging workflow includes mass spectrometry images of corynantheidine, GABA, and glutamine, fluorescence images of the μ-opioid receptors, fluorescence images of cell nuclei, and brightfield microscopy of the H&E stained tissue. Images visualized (**c**) prior to and (**d**) following image fusion highlight improvements in spatial sharpening and co-registration. (**e**) The areas highlighted in the white boxes were magnified.

Magnification of the mass spectrometry images allows for appreciation of the spatial sharpening enabled by image fusion (**Figure 4a-d**). The higher spatial resolution enabled by image fusion allows for the resolution of some nuclei in the predicted corynantheidine and glutamine mass spectrometry images (**Supplemental Figure 6**). IHC fluorescence images of the μ-opioid receptor were acquired on serial tissue sections to provide additional insight into alkaloid neuropharmacology (**Supplemental Figure 9**). Previous research has demonstrated the affinity of kratom alkaloids towards opioid receptors in cell-based assays.^45–47^ Visualizing the spatial relationship between the alkaloid and opioid receptors may provide insight into potential interactions of corynantheidine with these receptors after the alkaloid crosses the blood-brain barrier. Predicted image fusion results suggest localization of corynantheidine in the white matter. The white matter mainly consists of glial cells and myelinated axons.^56,57^ Corynantheidine shows a lower concentration in the glial cells and a higher concentration in the myelinated axons within the white matter (**Figure 4**). The glial and granular cells in the mass spectrometry image were identified using the cell nuclei visualized in the H&E image, which was acquired on the same tissue section as the mass spectrometry image. The μ-opioid receptor is mainly localized to the granular cells within the granular layer (**Supplemental Figure 9**). Low concentrations of the μ-opioid receptors located within the glial cells in the white matter are also observed. The glial and granular cells in the IHC fluorescence image were identified using the cell nuclei visualized in the DAPI fluorescence image, which was acquired on the same tissue section as the IHC fluorescence image (**Supplemental Figure 9**). Similar distributions of the μ-opioid receptors in the cerebellum of rat brain tissues have been previously reported.^58,59^ Comparing the alkaloid distribution to the μ-opioid receptor distribution, it appears that corynantheidine is present in lower concentrations in cells where receptors are located, which potentially implies that only small amounts of corynantheidine are penetrating cells where μ-opioid receptors are located. Permeability tests can be performed on *in vitro* cell monolayer models to test this hypothesis.^60^ High concentrations of corynantheidine are observed in the myelinated axons inside the white matter where the μ-opioid receptor is not observed, suggesting other receptors may be important in understanding the mechanism of action for kratom alkaloids. Image fusion predictions of GABA and glutamine do not allow for resolution of cell nuclei, suggesting the need for higher quality initial imaging mass spectrometry data for these analytes in order to provide better spatial sharpening predictions.

## DISCUSSION

In this study, we have described a multi-modal imaging workflow to study the neuropharmacology of the kratom alkaloid, corynantheidine. The multi-modal workflow was applied to brain tissue collected from a rat intravenously dosed with corynantheidine and consisted of mass spectrometry images (*i.e.*, of corynantheidine as well as of GABA and glutamine, two metabolites in the chronic pain pathway), an IHC fluorescence image of the μ- opioid receptor, and an H&E image showing tissue and cell morphology. Single-cell resolution is difficult to achieve using imaging mass spectrometry but is readily achieved with the other two imaging modalities. The mass spectrometry image resolution was improved roughly 10- fold through an image fusion workflow that models the relationship between the intensity values in the mass spectrometry images and the features extracted from a high-resolution H&E image.

Evaluation parameters are important criteria for determining the fidelities of the predicted images. The evaluation parameters proposed in this study can also be extended to other biological applications of image fusion. The accuracy of the predicted intensity values is evaluated through the relative average residual value, while the correlation scores evaluate how well the model is predicting the pattern of the image. Our results highlight that the models can have different degrees of spatial sharpening despite using the same input H&E image, suggesting the degree of spatial sharpening needs to be visually accessed, which has not been extensively described previously. Standard deviation images enable the user to determine whether the predictions in specific image regions are robust enough for the application at hand. Overall, the user can decide the most important evaluation criteria depending on the application, and different models will work better in different applications. Random forest regression and 2-D CNN models perform better than conventional linear and PLS regression models in predicting high spatial resolution mass spectrometry images for our application. Mass spectrometry and H&E microscopy images are generated through significantly different detection technologies. As such, a nonlinear regression model such as a random forest model may better describe the complex relationship between the intensity values in the mass spectrometry image and the manually extracted H&E image features compared to the linear models. The 2-D CNN model also performed well here, likely due to its ability to identify and extract features from the H&E image that are most important to defining the cross-modality relationship.

These two models enabled spatial sharpening of corynantheidine, GABA, and glutamine images from 25 μm to 2.5 μm. These higher spatial resolution images were then co-registered with both an IHC fluorescence image of the μ-opioid receptor and an H&E image. The abundance of corynantheidine appears lower in the glial and granular cells, which is where μ- opioid receptors are mainly localized and potentially implies limited penetration of the alkaloid into the cell bodies where μ-opioid receptors are located. This hypothesis can be tested in the future by performing permeability tests on *in vitro* cell monolayer models.^60^ While receptor co- localization alone is not sufficient to establish functional significance, these data can be used to formulate hypotheses for mechanisms of action for future studies into alkaloid metabolism. In general, image fusion enables more accurate integration of mass spectrometry and microscopy images for multi-modal imaging analyses compared to the original mass spectrometry images (**Figure 4**). This multi-modal imaging dataset is acquired on three serial sections that are separated by 10 μm each. Image fusion results in mass spectrometry images predicted to approximately 2.5 μm spatial resolution (*i.e.*, cellular to subcellular resolutions). However, the spacing of the serial tissue sections will result in spatial variations of single cells from section to section, which reduces accuracy and complicates image interpretation. Future analyses will develop sample preparation protocols that enable all necessary imaging modalities to be performed on the same tissue section.^61^ Alternatively, advanced registration methods, such as nonlinear stochastic embedding, can be employed to better account for differences in cell location.^50,62,63^ Despite these limitations, our results here still demonstrate the potential of multi-modal imaging to provide more comprehensive spatial information in pharmaceutical research. Future studies will also focus on developing a workflow to acquire all images on a single tissue section and on incorporating IHC fluorescence images of other receptors that potentially interact with alkaloids such as 𝜶-adrenoceptors and serotonin receptors. ^61,64–67^

## METHODS

### Materials

*Trans*-2-[3-(4-*tert*-Butylphenyl)-2-methyl-2-propenylidene]malononitrile (DCTB), 1,5- diaminonapthalene (DAN), γ-aminobutyric acid (GABA) standard, glutamine standards, Phloxine B, hematoxylin, sodium iodate, and aluminum potassium sulfate dodecahydrate were purchased from Sigma-Aldrich (St. Louis, MO, USA). Corynantheidine standard was extracted from dried *Mitragynine speciosa* leaves (Pure Land Ethnobotanicals, Madison WI, USA) using methanol or ethanol, and then fractioned and dried using acid-base extraction and column chromatography. Proton nuclear magnetic resonance (^1^H NMR), carbon nuclear magnetic resonance (^13^C NMR), high-performance liquid chromatography-photometric diode array (HPLC-PDA), and ultra-performance LC-quadrupole time of flight (UPLC-QTOF) were used to ensure the purity of the standard were 98% or above.^12^ Epredia™ Cytoseal™ mountant XYL, glycerol, Eosin Y intensified solution, μ-opioid receptor polyclonal antibody, donkey anti-rabbit IgG (H+L) highly cross-absorbed secondary antibody (Alexa fluor plus 488, absorption at 497nm, emission at 517nm), and ProLong^TM^ Gold antifade mountant with DAPI were purchased from ThermoFisher Scientific (Waltham, MA). Xylene was purchased from Honeywell International Inc. (Charlotte, NC).

### Imaging mass spectrometry

Sprague Dawley rats were intravenously injected with corynantheidine and were sacrificed 30 minutes after dosage. The brains were immediately removed, rinsed with saline, and stored at -80°C. 10 μm thick sagittal cross-sections of the dosed brains were collected using a Leica CM3050S Research Cryostat (Lecia Biosystems, Wetzlar, Germany). The chamber temperature was set to -24°C and the object temperature was set to -22°C during sectioning. Sections were mounted onto indium tin oxide (ITO)-coated glass slides (Delta Technologies, Loveland, CO, USA). Either DAN (targeting GABA and glutamine) or DCTB (targeting corynantheidine) matrix was applied using a home-built sublimation apparatus (DAN: 9 minutes at 115°C and < 70 mTorr, resulting in 1-2 mg of matrix deposited on the slide; DCTB:8.5 minutes at 100°C and <70 mTorr, resulting in 1-2 mg of matrix deposited on the slide). All imaging experiments were performed on a 7T solariX Fourier transform ion cyclotron resonance (FT-ICR) mass spectrometer (Bruker Daltonics, Billerica, MA) equipped with an Apollo II dual MADLI/ESI source. The laser for the MALDI source is a Smartbeam II Nd:YAG laser system (2kHz, 355nm). Images were acquired at 25 μm and 75 μm spatial resolution in both positive (*i.e.*, for corynantheidine analysis) and negative ion mode (*i.e.*, for GABA and glutamine analysis) using 300 laser shots and 20% global laser power for the corynantheidine images and 500 laser shots and 28% global laser power for the GABA and glutamine images. The smartwalk was set to the same value as the spatial resolution. Data were collected from *m/z* 100 to 1,000 using a 0.2447 s time-domain transient length, which resulted in a mass resolving power of ∼29,500 at *m/z* 367. Continuous accumulation of selected ions (CASI) was used to isolate and enrich ions of interest (10 Da wide, centered at the nominal mass of the target ion).^13,68^ Identification of corynantheidine, GABA, and glutamine was confirmed via comparisons to reference standards and via high-resolution accurate mass measurements with mass accuracy errors less than 5 ppm.

### Tissue Staining

H&E staining was performed by placing the slide in 95% ethanol, 70% ethanol and Milli-Q water sequentially each for 30 seconds. The slide was then placed in hematoxylin solution (1 g of hematoxylin, 45.93 g of aluminum potassium sulfate dodecahydrate, and 0.1 g of sodium iodate dissolved in 379 mL of Milli-Q water and 100 mL of glycerol) for 2 minutes. Next, the slide was sequentially placed in solutions of Milli-Q water, 70% ethanol, 95% ethanol, and eosin (1 g of phloxine B and 50 mL of Eosin Y intensified solution dissolved in 100 mL of Milli-Q water) for 20 seconds, 30 seconds, 30 seconds, and 1 minute, respectively. Finally, the slide was placed sequentially in 95% ethanol, 100% ethanol, and Xylene for 30 seconds, 30 seconds, and 2 minutes, respectively. A slide mounting process is commonly performed following H&E staining to prevent loss or oxidation of the stain due to contact with the air. However, a coverslip was not mounted to the slide in our workflow to maximize color contrast (**Supplemental Figure 10**).

IHC was performed to visualize μ-opioid receptors. Briefly, two dosed tissue sections were mounted on a microscope slide. One section was used as the control section to verify the absence of nonspecific binding. Tissue sections were covered with 4% paraformaldehyde solution and incubated in a 4°C freezer for 15 minutes and then rinsed with PBS buffer (7.4 PH). The slide was incubated in a Coplin jar filled with blocking solution (3% bovine serum albumin (BSA) in PBS buffer) for 1 hour at room temperature. Following incubation, the non-control section was exposed to the primary antibody solution (μ-opioid receptor polyclonal antibody, 1:300 dilution with 1% BSA and 0.3%TritonX-100 in PBS buffer) and incubated in 4°C freezer for roughly 19 hours. The slide was then washed with PBS buffer and both tissue sections were exposed to the secondary antibody solution (donkey anti-rabbit IgG (H+L) highly cross-absorbed secondary antibody, 1:400 dilution with 1% BSA and 0.3%TritonX-100 in PBS buffer) and incubated at room temperature for 60 minutes. The secondary antibody is tagged with an Alex fluor plus fluorophore. The slides were washed with PBS solution and then covered with the ProLong^TM^ Gold antifade mountant with DAPI as a nuclear counter stain and sealed with a coverslip. A control experiment omitting the primary antibody was performed on a serial tissue section to verify the absence of nonspecific binding (**Supplemental Figure 11**).

### Image fusion

The image fusion workflow used in this study is based on the method reported by Van de Plas, *et al.*^41^ The workflow can be divided into two main steps: (1) accurate co-registration of the mass spectrometry image with the H&E microscopy image and (2) modeling the cross-modality relationship between the mass spectrometry data and the features of the H&E image (**Figure 1**). MALDI imaging mass spectrometry is first performed on the target tissue. An autofluorescence image of the tissue following imaging mass spectrometry is then acquired using the absorption/emission profile of DAPI (absorption at 358 nm, and emission at 465 nm) with the matrix still present on the tissue. The autofluorescence image clearly reveals the laser ablation marks in the matrix coating produced during MALDI imaging analysis.^50^ The matrix is then washed off and stained via hematoxylin and eosin (H&E) and scanned to a TIF file format via brightfield microscopy using an upright microscope (Imager M2, Zeiss Axio). The H&E microscopy image is compressed so that the number of pixels in the H&E image is approximately 100x the number of pixels in the mass spectrometry image (*i.e.*, the H&E brightfield microscopy image will be 10x higher in spatial resolution compared to the mass spectrometry image).

The mass spectrometry image was aligned to the H&E image using a workflow similar to that proposed by Patterson, *et al.* and was performed manually in MATLAB.^46^ Briefly, the mass spectrometry image is first registered to the autofluorescence image by matching 3-4 pixels in the mass spectrometry image to their corresponding laser ablation marks shown in the autofluorescence image. The autofluorescence image is then also registered to the H&E image by finding 3-4 corresponding tissue structure points in both images. This connects the coordinate system of the mass spectrometry image to that of the H&E image through the autofluorescence image and thus allows for registration of the mass spectrometry image to the H&E image (**Figure 1**). Non affine transformations are used for all transformations because all images are acquired from the same tissue region.

Cross-modality model construction begins by expanding the three RGB features of the H&E image to 120 features using the rgb2lab, rgb2hsv, rgb2ntsc, rgb2ycbcr, rangefilt, and entropyfilt functions in MATLAB (**Supplemental Table 1**). Models including linear regression (MATLAB), partial least squares regression (MATLAB), random forest regression (using the DecisionTreeRegressor function from the sklearn library Python), and two-dimensional convolutional neural networks (using the keras library in Python) are used to describe the relationship between the intensity of the target analyte in the mass spectrometry image and the extracted features of the H&E image. The model predicts the higher spatial resolution mass spectrometry image by using the H&E image features as the input data. The reported spatial resolution of the predicted mass spectrometry images was equal to the spatial resolution of the H&E image used as the input. Evaluations of the models were performed via calculation of the correlation score, calculation of the relative average residual (**Equation 1**), and visual verification of the spatially sharpened predicted image. The correlation score is the R^2^ value obtained by plotting the intensity value for each pixel in the predicted image against the corresponding intensity value in the original mass spectrometry image using the plotregression function in MATLAB. Residuals are calculated by:

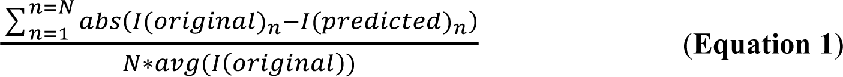

where N is the total number of pixels, I(original)n is the intensity value of the n^th^ pixel in the original mass spectrometry image, and I(predicted)n is the intensity value of the n^th^ pixel in the predicted mass spectrometry image. The violin plots of the relative residual values used to calculated the relative average residual were generated using the seaborn package in Python. The regression plots of the original mass spectrometry image intensity values versus the predicted mass spectrometry image intensity values were generated using the plotregrssion function in MATLB.

### Standard deviation images

The 10 bootstrap models for the random forest regression models were generated by performing random resampling with replacement 10 times using an in-house Python script. The random forest model was then trained with the bootstrap dataset and then applied to the original dataset. Random resampling is not applicable to the 2-D CNN models due to the spatial awareness of these models. 294*294 blocks (around 10% of the total pixel number) were removed from the dataset to generate the bootstrap dataset. Only 5 bootstrap models were trained due to the long training times of the CNN models. The trained models were then applied to the original dataset (**Supplemental Figure 7**).

## CODE AVAILABILITY

Source code is publicly available on GitHub: https://github.com/Prentice-lab-UF/Image-fusion-

## ACKNOWLEDGEMENTS

This work was supported by the National Institutes of Health (NIH) under award R01 GM138660 (National Institute of General Medical Sciences [NIGMS]) (BMP), by a contribution from Eli Lilly and Company (BMP, under award R01 DA047855 (National Institute of Drug Abuse (NIDA]) (CRM), and under award UG3/UH3 DA048353 (NIDA) (CRM). The authors would like to thank Prof. Raf Van de Plas at the Delft University of Technology (TU Delft) in the Netherlands for helpful guidance and discussions throughout this project.

