## Supplemental Information for "A multi-modal image fusion workflow incorporating MALDI imaging mass spectrometry and microscopy for the study of small pharmaceutical compounds"

**Table of contents:**

**S3** Supplemental Table 1: MATLAB functions used to extract H&E image features
**S4** Supplemental Table 2: Structure of the 2D-CNN model
**S5** Supplemental Table 3: PLS regression and random forest regression model optimization for corynantheidine
**S6** Supplemental Table 4: PLS regression and random forest regression model optimization for GABA
**S7** Supplemental Figure 1: Overlay of 75 µm mass spectrometry images and H&E images
**S8** Supplemental Figure 2: Overlay of 25 µm mass spectrometry images and H&E images
**S9** Supplemental Figure 3: Image fusion model optimization for mass spectrometry image of glutamine acquired at 75 µm spatial resolution
**S10** Supplemental Figure 4: Zoomed in predicted images acquired from 75-µm mass spectrometry images
**S11** Supplemental Figure 5: Zoomed in predicted images acquired from 25-µm mass spectrometry images
**S12** Supplemental Figure 6: Zoomed in predicted images of corynantheidine and glutamine used in the final multi-modal workflow
**S13** Supplemental Figure 7: Bootstrap models for 2-D CNN models
**S14** Supplemental Figure 8: Standard deviation images
**S15** Supplemental Figure 9: Zoomed in Fluorescence images of the µ-opioid receptors and cell nuclei
**S16** Supplemental Figure 10: Brightfield microscopy image of H&E-stained tissue
**S17** Supplemental Figure 11: Fluorescence images

**Supplemental Table 1**. MATLAB functions are used to extract 120 features from H&E image.

| Feature number | Feature/feature generating function |
|---|---|
| 1-3 | RGB value |
| 4-6 | rgb2lab |
| 7-9 | rgb2hsv |
| 10-12 | rgb2ntsc |
| 13-15 | rgb2ycbcr |
| 16-30 | PCA components |
| 31-60 | entropyfilt |
| 61-120 | rangefilt |

**Supplemental Table 2:** The structure of the neural network layers is described based on the parameters and activation. The “means_sqaured_error” loss function and “adam” optimizer was used. The epoch number was set to 10,000.

| Layer | Type | Parameters | Activation |
|---|---|---|---|
| 1 | BatchNormalization | NA | NA |
| 2 | ZeroPadding2D | Padding (1,1) | NA |
| 3 | Conv2D | 64,kernel_size (3,3),kernel_initializer (normal) | ReLu |
| 4 | ZeroPadding2D | Padding (1,1) | NA |
| 5 | Conv2D | 32,kernel_size (3,3),kernel_initializer (normal) | ReLu |
| 6 | ZeroPadding2D | Padding (1,1) | NA |
| 7 | Conv2D | 1,kernel_size (1,1) | NA |

**Supplementary Table 3.** PLS regression and random forest regression were optimized for image fusion performed using 75 μm spatial resolution mass spectrometry images of corynantheidine. PLS regression was optimized in terms of the component number. Random forest regression was optimized in terms of the number of H&E image features used as the input and the number of decision trees included. Optimization was based on the relative average residual value and correlation score ($R^2$).

| **PLS components** | **Relative average residual** | **$R^2$** |
|---|---|---|
| 35 | 0.380 | 0.789 |
| 37 | 0.380 | 0.790 |
| 40 | 0.380 | 0.791 |
| 45 | 0.384 | 0.789 |
| 50 | 0.406 | 0.779 |
| **Number of features_number of trees** | **Relative average residual** | **$R^2$** |
| 3_1 | 0.335 | 0.799 |
| 3_10 | 0.321 | 0.830 |
| 3_100 | 0.321 | 0.834 |
| 15_1 | 0.334 | 0.801 |
| 15_10 | 0.320 | 0.831 |
| 15_100 | 0.318 | 0.834 |
| 30_1 | 0.334 | 0.800 |
| 30_10 | 0.320 | 0.831 |
| 30_100 | 0.318 | 0.834 |
| 60_1 | 0.186 | 0.880 |
| 60_10 | 0.137 | 0.972 |
| 60_100 | 0.119 | 0.983 |
| 120_1 | 0.178 | 0.890 |
| 120_10 | 0.130 | 0.975 |
| 120_100 | 0.113 | 0.984 |

**Supplementary Table 4.** PLS regression and random forest regression were optimized for image fusion performed using 75 μm spatial resolution mass spectrometry images of GABA . PLS regression was optimized in terms of the component number. Random forest regression was optimized in terms of the number of H&E image features used as the input and the number of decision trees included. Optimization was based on the relative average residual value and correlation score ($R^2$).

| PLS components | Relative average residual | $R^2$ |
|---|---|---|
| 5 | 0.171 | 0.775 |
| 8 | 0.170 | 0.771 |
| 10 | 0.176 | 0.751 |
| 11 | 0.176 | 0.737 |
| 15 | 0.224 | 0.622 |
| **Number of features_number of trees** | **Relative average residual** | **$R^2$** |
| 3_1 | 0.154 | 0.802 |
| 3_10 | 0.146 | 0.832 |
| 3_100 | 0.145 | 0.835 |
| 15_1 | 0.151 | 0.803 |
| 15_10 | 0.146 | 0.832 |
| 15_100 | 0.145 | 0.835 |
| 30_1 | 0.151 | 0.802 |
| 30_10 | 0.146 | 0.832 |
| 30_100 | 0.144 | 0.835 |
| 60_1 | 0.079 | 0.883 |
| 60_10 | 0.059 | 0.972 |
| 60_100 | 0.053 | 0.982 |
| 120_1 | 0.076 | 0.891 |
| 120_10 | 0.057 | 0.974 |
| 120_100 | 0.050 | 0.984 |

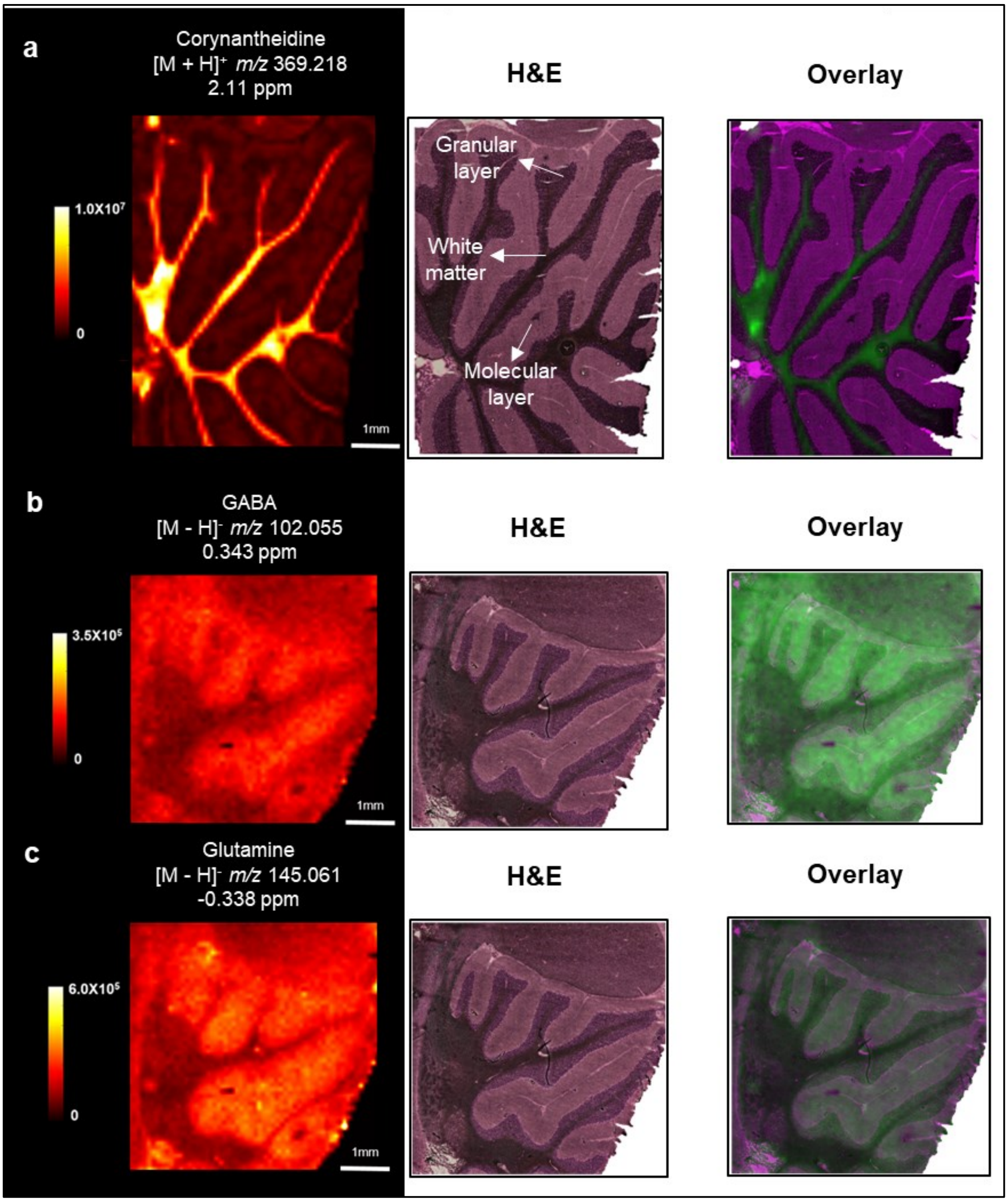


**Supplemental Figure 1.** Imaging mass spectrometry analyses acquired at 75 µm spatial resolution targeting (**a**) corynantheidine, (**b**) GABA, and (**c**) glutamine are shown alongside H&E brightfield microscopy images of the same tissue section. H&E images were acquired without a coverslip to maximize contrast. Brain tissue is derived from a Sprague Dawley rat dosed with corynantheidine. Image mass spectrometry is co-registered with microscopy, allowing for tissue overlays (purple: H&E image, green: mass spectrometry image).

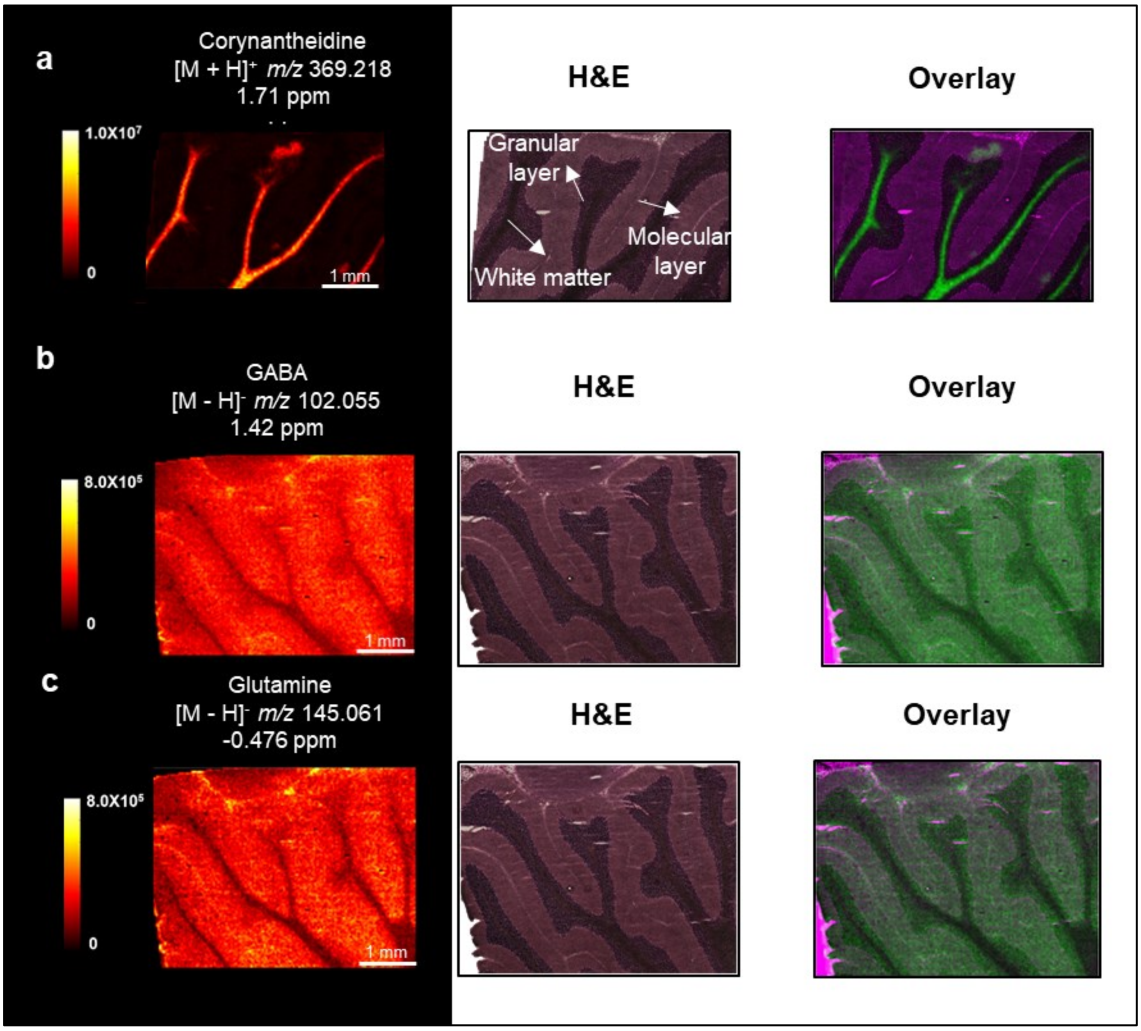


**Supplemental Figure 2.** Imaging mass spectrometry analyses acquired at 25 μm spatial resolution targeting (**a**) corynantheidine, (**b**) GABA, (**c**) and glutamine are shown alongside H&E brightfield microscopy images of the same tissue section. H&E images were acquired without a coverslip to maximize contrast. Brain tissue is derived from a Sprague Dawley rat dosed with corynantheidine. Image mass spectrometry is co-registered with microscopy, allowing for tissue overlays (purple: H&E image, green: mass spectrometry image).

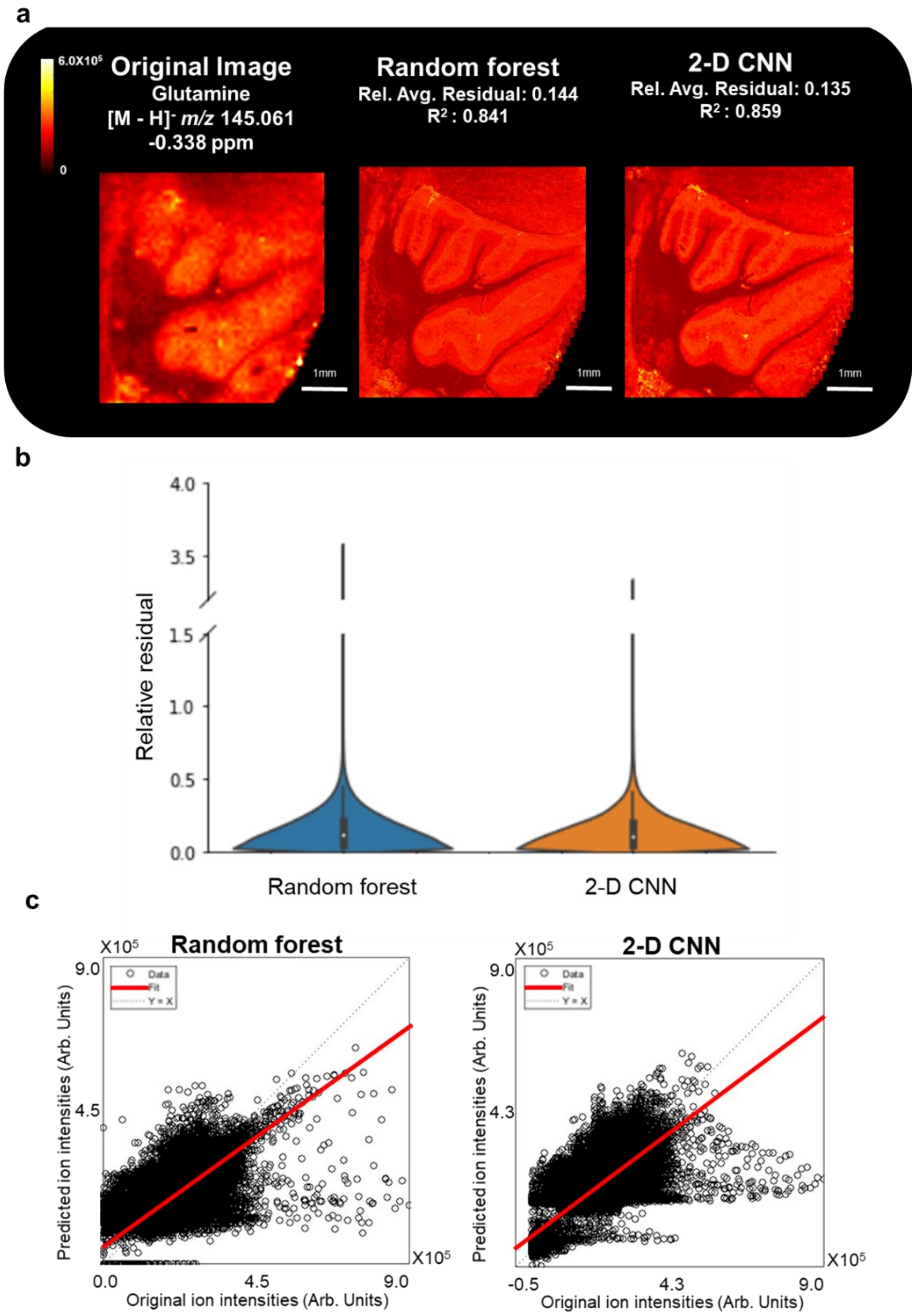


**Supplemental Figure 3.** (**a**) Image fusion is performed on 75 µm spatial resolution mass spectrometry images of glutamine using random forest regression and 2-D CNN. The relative average residual (Rel. Avg. Residual) and correlation score ($R^2$) are calculated for each predicted mass spectrometry image. (**b**) The relative residual value for each pixel, which was utilized in calculating the relative average residual, is shown using a violin plot. (**c**) The original mass spectrometry image intensity values are plotted against the predicted mass spectrometry image intensity values to acquire the correlation score.

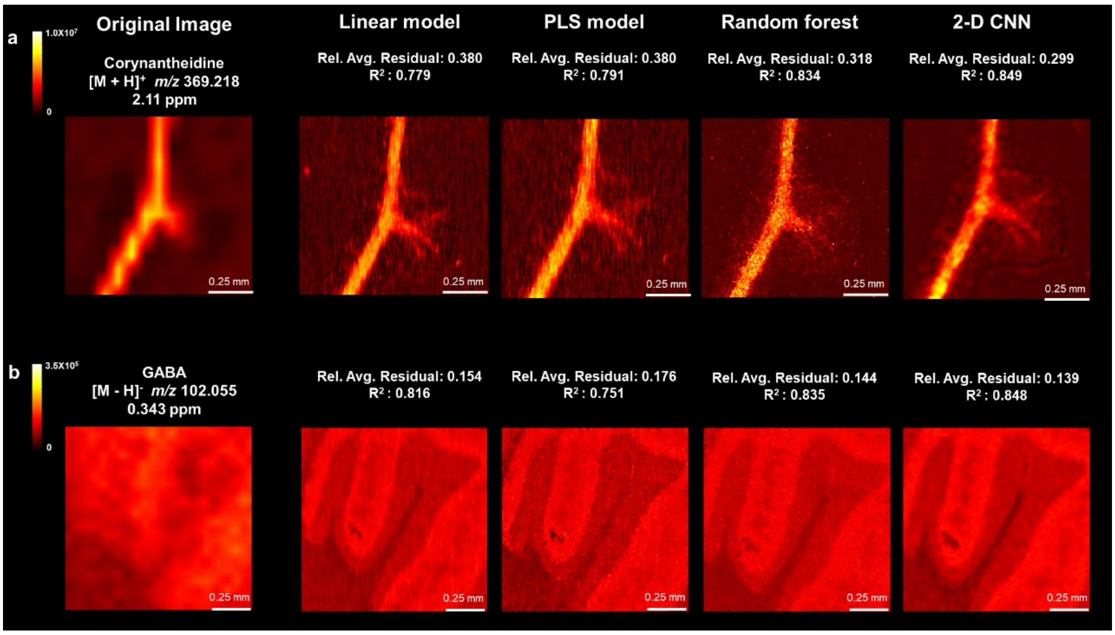


**Supplemental Figure 4.** Image fusion is performed on mass spectrometry images of (**a**) corynantheidine and (**b**) GABA acquired at 75 µm spatial resolution using linear regression, PLS regression, random forest regression, and 2-D CNN. The predicted images are magnified to show the varying levels of spatial sharpening for each model.

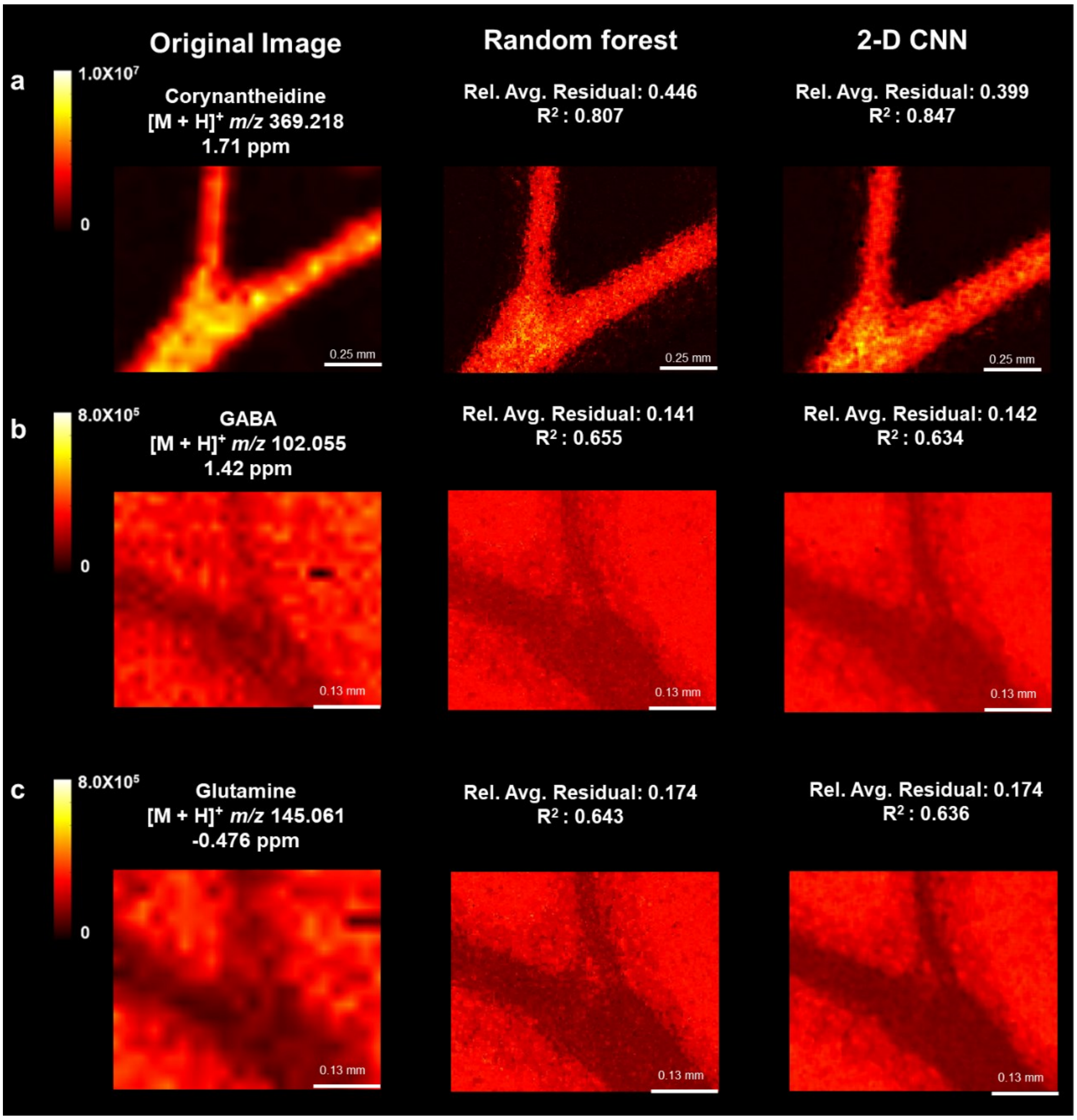


**Supplemental Figure 5.** Image fusion performed using 25 μm spatial resolution mass spectrometry images of (**a**) corynantheidine, (**b**) GABA, and (**c**) glutamine using random forest regression and 2-D CNN models results in predicted 2.5 μm spatial resolution images. The predicted images are magnified to show the varying levels of spatial sharpening for each model.

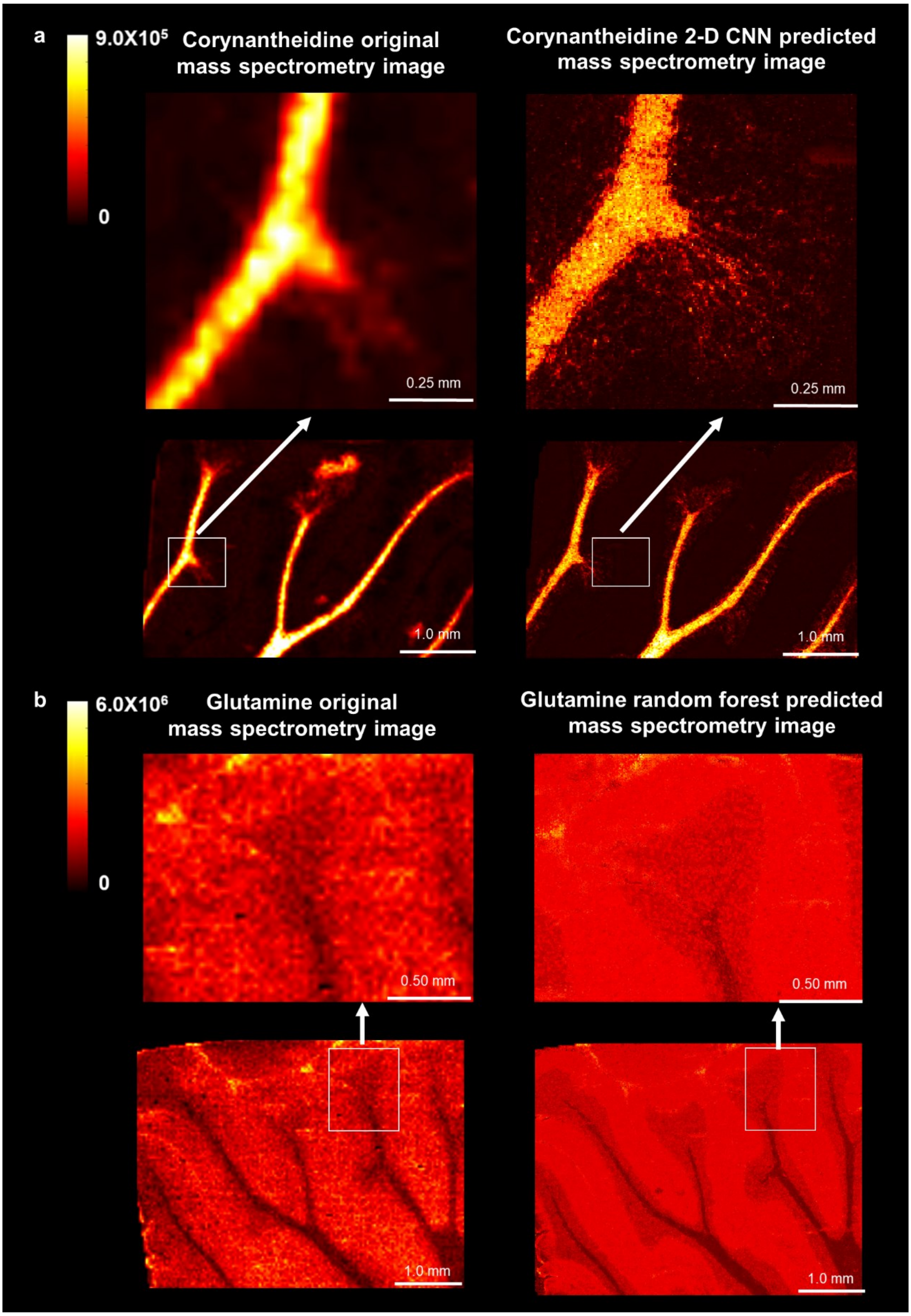


**Supplemental Figure 6.** Optimized image fusion models are used to predict 25 µm spatial resolution mass spectrometry images of (**a**) corynantheidine and (**b**) glutamine to 2.5 µm spatial resolution.

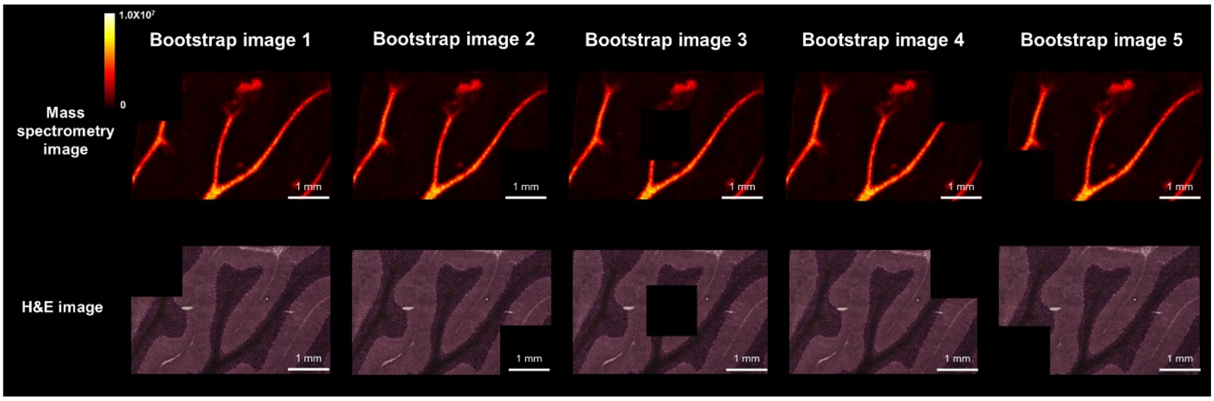


**Supplemental Figure 7.** Bootstrap models are created for the 2-D CNN models by removing 294*294 blocks from the dataset. H&E images were acquired without a coverslip to maximize contrast.

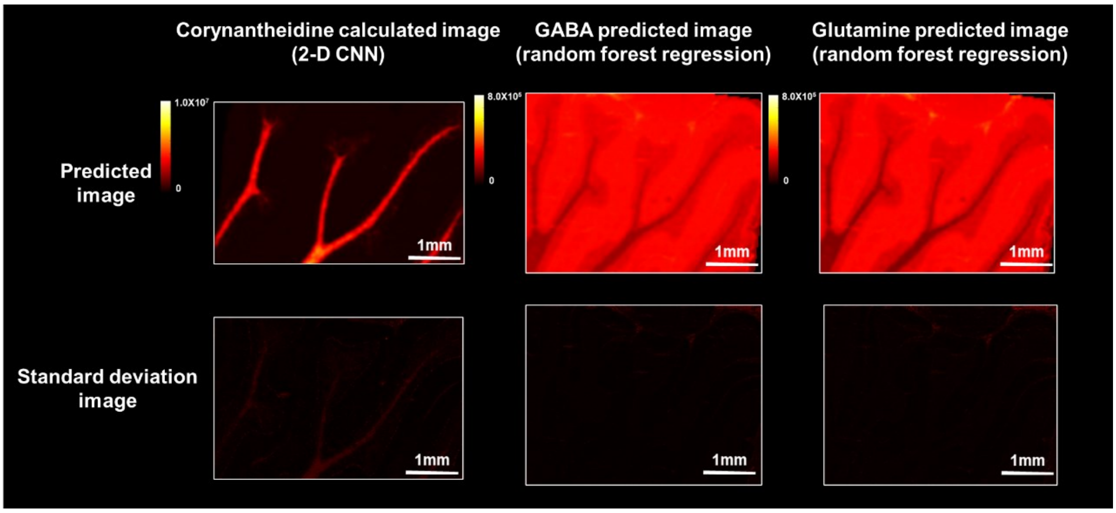


**Supplemental Figure 8**. Standard deviation images are calculated for the 2D-CNN model used for the corynantheidine mass spectrometry image and the random forest model used for the GABA and glutamine mass spectrometry images.

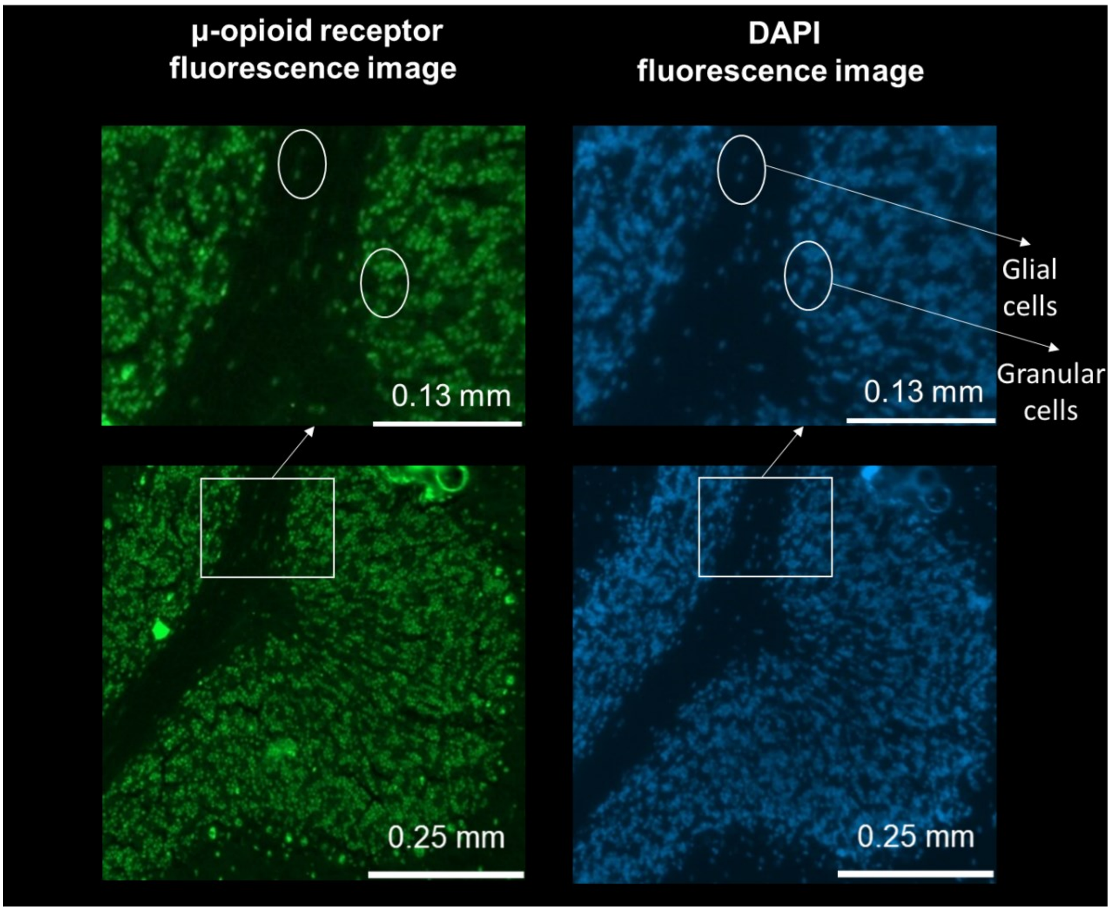


**Supplemental Figure 9**. Fluorescence images of μ-opioid receptors and cell nuclei (identified using DAPI staining) are magnified to visualize the localization of the μ-opioid receptors in the glial and granular cells (highlighted using white circles).

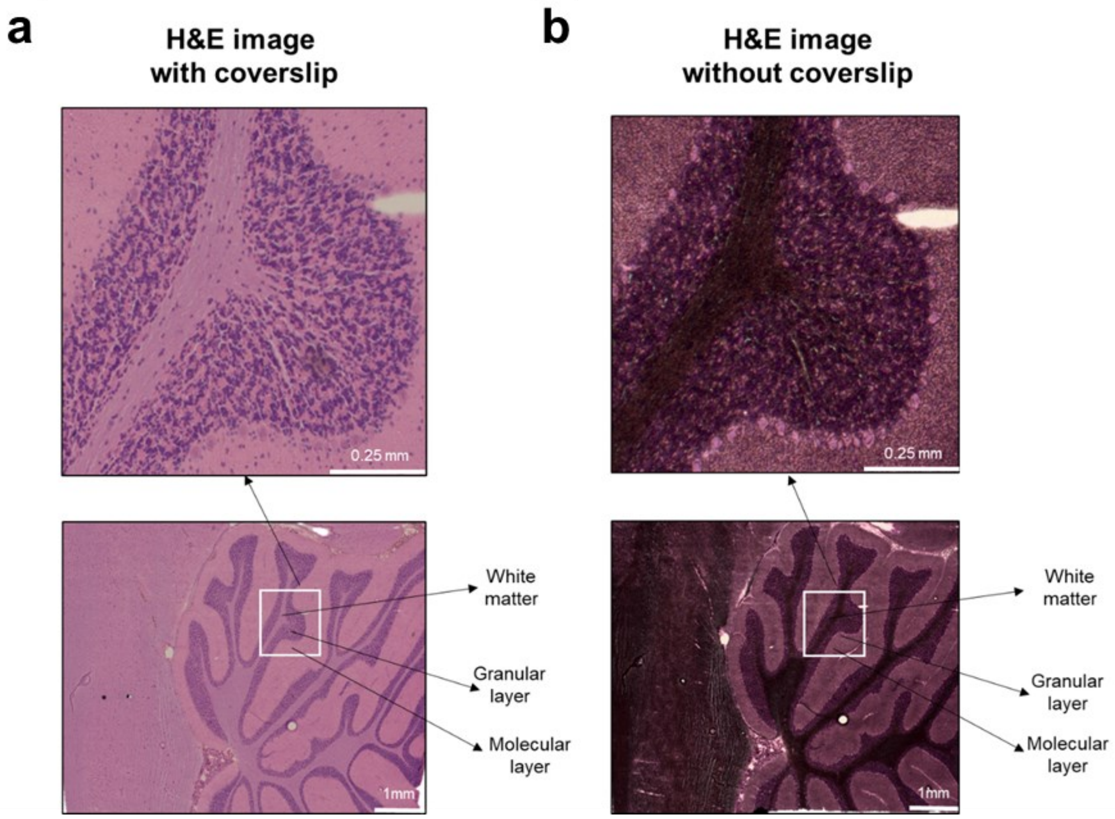


**Supplemental Figure 10.** Brightfield microscopy images are acquired (**a**) with and (**b**) without a coverslip for H&E-stained brain tissue sections from a Sprague Dawley rat dosed with corynantheidine.

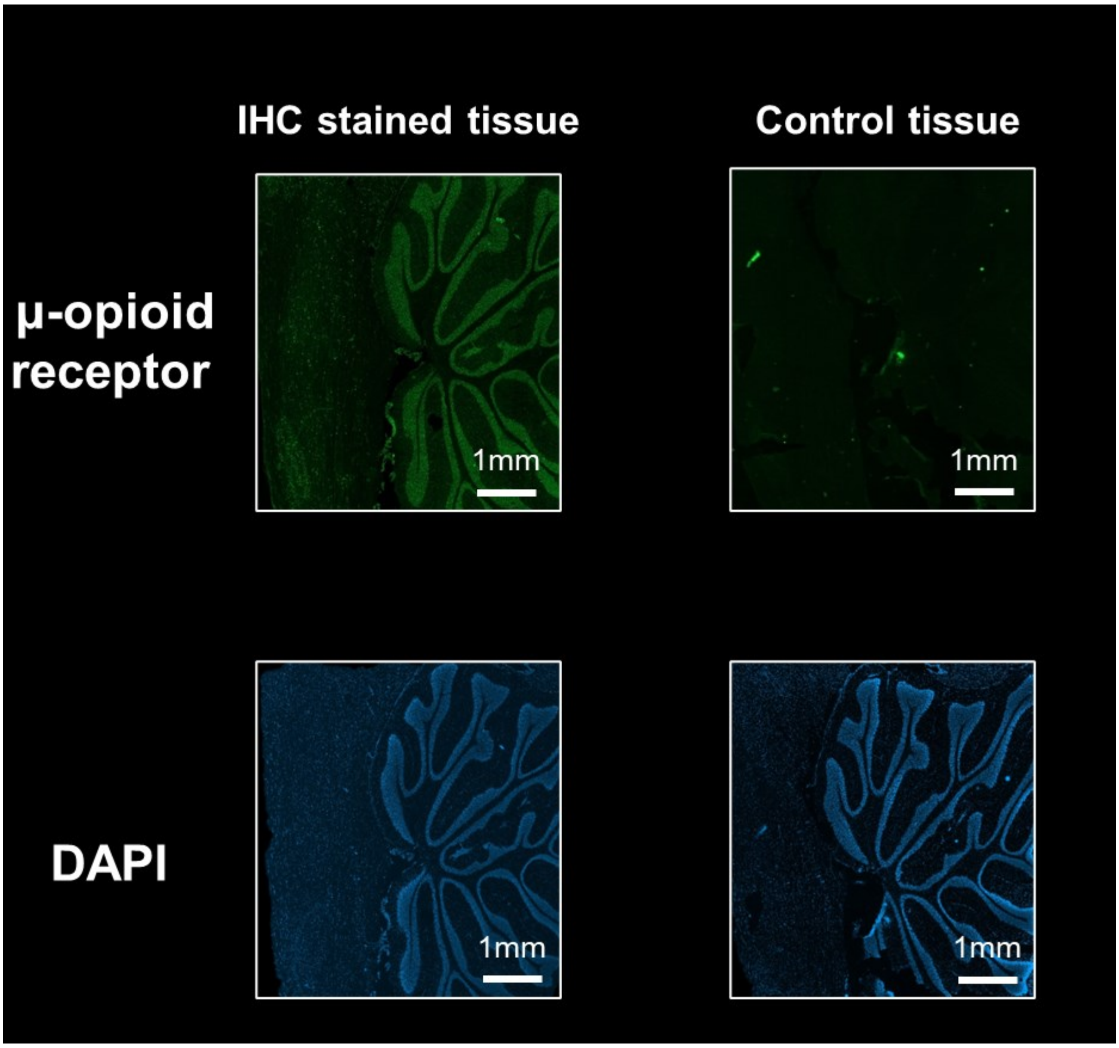


**Supplemental Figure 11**. The µ-opioid receptors are first labeled with a primary antibody, which is subsequently targeted with a secondary antibody tagged to an Alexa fluor plus fluorophore. A control experiment omitting the primary antibody performed on a serial section verifies the absence of nonspecific binding. Fluorescence DAPI images enable visualization of the cell nucleus.
